# Detection of adaptation to light environment in myopia-associated loci Diversity of the myopia genetic background

**DOI:** 10.1101/2022.10.27.513974

**Authors:** Tian Xia, Kazuhiro Nakayama

**Author notes:** The authors have declared that no competing interests exist.

## Abstract

The time spent outdoors and exposure to bright sunlight have been reported to play important roles in preventing myopia progression. Myopia prevalence is generally low in Europe, and we hypothesized that local adaptation of ancestors in insufficient sunlight regions might underlie this low prevalence in Europeans. To verify this conjecture and understand how ancestry adaptation influenced on the diversity of myopia genetic background, polygenic risk scores (PRS) was calculated and revealed that the genetic risk of myopia increased as latitude decreased in the 1000 genome projects (1KGP) EUR, and we further determined selection signatures near the rhodopsin (*RHO*) coding region in 1KGP Finnish. The derived allele frequencies of identified loci correlated with sunshine duration worldwide, and the allele age was estimated to be ∼20,000 years, coincidently after the divergence of Europeans and East Asians. *TSPAN10* (rs9747347) is significantly associated with both myopia and pigmentation in Europeans, and we found selection favored the myopia risk allele in EUR. Haplotype comparison highlighted the divergence between EAS and EUR in *RHO* and pigmentation-associated loci, and the frequency diversity in Admixed American (AMR) of these loci was found to originate from recent admixture and local adaptation. We concluded that local adaptation played a role in myopia prevalence deviation, and selective pressure in the myopia-associated loci may originate from adaptation to sunlight environments rather than myopia itself. These findings reveal traces of evolution history of myopia’s genetic background, providing a new aspect of understanding the patterns and diversity of myopia prevalence.

**Author Summary:** The explanation for the distinct bias of myopia prevalence among ethnicities remains controversial, we took an evolutionary approach to understanding the history and diversity of the genetic background of myopia. We detected selection signatures of myopia-associated loci near the rhodopsin gene (*RHO*) in the 1000 Genome Project Europeans. Ambient light exposure is crucial for myopia; *RHO* is mainly expressed in rod cells and is extremely sensitive to light, indicating that local adaptation might contribute to myopia prevalence bias. Parallel geographical disposition of allele frequencies of myopia-associated loci in *RHO* and pigmentation-associated markers was observed, implying that selective pressure in myopia-associated loci may originate from adaptation events involved in sunlight exposure rather than myopia itself.

## Introduction

Myopia is reported as the most common cause of distance vision impairment globally[1], has emerged in the last century[2,3], and is commonly recognized as being caused by gene-environment interactions[4,5]. Time spent outdoors and under bright sunlight has been reported to play important roles in preventing myopia progression[6,7,8]. In addition, light intensity[9,10] has been reported to impact myopia progression; spectrum composition[11,12,13,14,15] has been demonstrated to be critical in animal models of refractive error; and spatial frequency spectra of the man-made environment was previously reported as a potential factor of the myopia epidemic[16].

Local adaptation to light environment has contributed to genetic and phenotypic differentiation among human populations, such as pigmentation[17]. Since vision and the eyes are products of evolution[18], a proportion of myopic genetic background divergence may originate from local adaptation. The continuous adaptation to various light environments could have even started right after modern humans migrated out of Africa ∼135 to 37 kya[19].

Regarding local adaptation, sunshine duration was more reliable than latitude in terms of light environmental factors. Sunshine exposure duration depends on latitude and cloud amount[20]. Even though the solar spectrum varies across latitudes due to the curvature of the Earth, clouds were found to have a greater influence on the solar spectrum than latitude. Clouds blocking the sun causes a higher proportion of blue in visible light[21,22], meanwhile the long-wavelength red light was recently found can slow the progression of myopia in children[23,24]. Moreover, sunshine duration refers to sunny hours during the daytime. People living in some extreme high-latitude regions are usually not exposed to sunlight during resting night hours during the midnight sun seasons.

As bright sunlight exposure is critical for myopia pregression, we hypothesize that people living in regions of less sunshine duration for thousands of years were capable of adapting to the modern indoor-centric environment, and eventually resulting in lower myopia prevalence. Besides some Northern European myopia studies 25, 26, 27, the prevalence distribution by continent1,28 may support this conjecture (Fig 1A, Table 1). To date, major large-scale myopia studies are European ancestry-centric29, 30, and sunshine duration varies drastically in Europe31, making Europeans the ideal subjects for this study.

**Table 1.** Frequencies and sunshine duration of rs2855558 derived allele (A).

| Population | Derived allele frequency | City of Sunshine Duration observation | Sunshine Duration(h/y) |
| --- | --- | --- | --- |
| FIN | 92.93% | Helsinki, Finland | 1,776 |
| Estonia | 90.96% | Tallinn, Estonia | 1,755 |
| North Sweden | 89.30% | Haparanda, Sweden | 1,770 |
| GBR | 84.78% | London, UK | 1,569 |
| Netherlands | 83.50% | Utrecht, Netherlands | 1,472 |
| TSI | 79.62% | Pisa, Italy | 2,269 |
| IBS | 77.57% | Madrid, Spain | 2,673 |
| MXL | 76.86% | Mexico City, Mexico | 2,217 |
| Qatari | 71.80% | Doha, Qatari | 3,431 |
| CLM | 70.21% | Barrancabermeja, Colombia | 2,169 |
| PUR | 68.10% | San Juan, Puerto Rico | 2,960 |
| GIH | 63.68% | Ahmedabad, India | 3,026 |
| PJL | 61.98% | Lahore, Pakistan | 3,038 |
| STU | 61.65% | Colombo, Sri Lanka | 2,625 |
| ITU | 60.19% | Bangalore, India | 2,369 |
| BEB | 53.49% | Calcutta, India | 2,114 |
| Korean | 45.91% | Seoul, South Korea | 2,108 |
| JPT | 42.79% | Tokyo, Japan | 1,813 |
| CHB | 42.23% | Beijing, China | 2,743 |
| CHS | 32.87% | Nanchang, China | 1,861 |
| CDX | 30.30% | Kunming, China | 2,336 |
| ACB | 11.46% | Guadeloupe, Caribbean islands | 2,768 |
| LWK | 7.92% | Kitale, Kenya | 2,603 |
| ESN | 4.54% | Enugu, Nigeria | 2,035 |
| MSL | 3.53% | Bo, Sierra Leone | 1,937 |
| GWD | 3.10% | Kolda, Senegal | 2,890 |
| YRI | 1.85% | Ironin, Nigeria | 2,284 |

Sunshine duration data (hours per year) are from the Hong Kong Observatory, allele frequencies are from NCBI, and target data.

**Fig 1.**
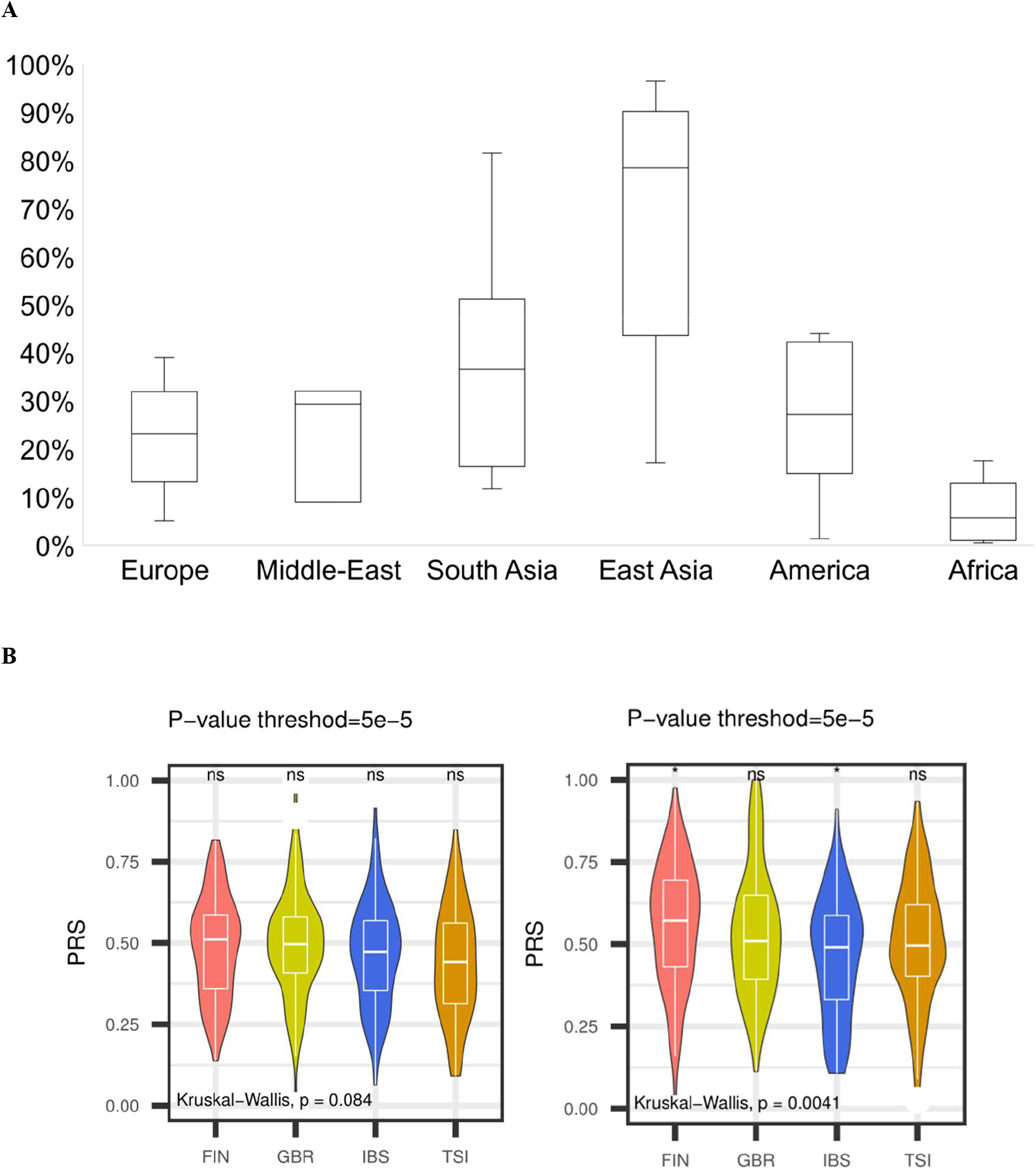
Global myopia prevalence and PRS distribution in 1KGP EUR. (A) Global myopia prevalence by region. Myopia prevalence records were extracted from previously studies[1,29,30], and re-arranged by continents, comprising 175 studies and 64 countries. (B) Distribution of the two second PRS settings at *P* <5.0× 10^−5^ by percentile; Left: PRS of 46,423 common scored variants in the first round. Right: PRS of 569,973 whole scored variants in the first round. Each violin plot represents the relative scores of the four populations. FIN: Finnish in Finland; GBR: British from England and Scotland; IBS: Iberian populations in Spain; TSI: Toscani in Italy. The lower four P thresholds all represent the myopia genetic risk that arises as latitude drops in FIN (60∼70°N), GBR (49∼59°N), IBS (27∼44°N), and TSI (35∼47°N).

This study aims to validate the conjecture that adaptation to light environment played a role in the divergence of myopia prevalence, and understand how ancestry adaptation influenced on the diversity of myopia genetic background. We compared polygenic risk scores (PRS)[32] distribution among 1KGP EUR populations, and detected selection signatures in myopia-associated loci using population branch statistics (PBS)[33] and haplotype-based statistics (*nSL*[34] and XP-*nSL*[35]). Although it is difficult to determine to what extend the genetic background variation dedicates to the global myopia prevalence bias, this study provided a new aspect of understanding the prevalence divergence and we call for flexible myopia prevention methods to apt different populations.

## Results

### PRS in 1KGP Europeans showing myopia risk predisposition correlated with latitude

The base and target data in the PRS were extracted from the European ancestry myopia GWAS summary statistics[29] and genotypes of the 1KGP populations[36], respectively. The ancestry composition in the base data (*N*=276,065) was 76.46% from UK Biobank (UKBB), 12.67% from the Genetic Epidemiology Research on Adult Health and Aging (GERA), and 10.86% from the Consortium for Refractive Error and Myopia (CREAM) participants[29]. To avoid over-fit for GBR and sampling bias during variant clumping due to general genetic distance (*e*.*g*., FIN and IBS), a two-round PRS analysis strategy was carried out in this study (*see Materials and Methods*).

PRS were calculated per individual at eight *P* thresholds (0.1, 0.05, 0.01, 5.0 × 10^−3^, 5.0 × 10^−5^, 5.0 × 10^−8^, 5.0 × 10^−12^ and 5.0 × 10^−20^) in 1KGP FIN(*N*=96), GBR(*N*=91), IBS(*N*=100), and TSI(*N*=107). Although the best-fit *P* threshold cannot be anchored without the phenotype of the subjects, the relative PRS orders at strata of *P* thresholds still present a general genetic risk predisposition. In the first PRS round, myopia genetic risk increased as the latitude decreased in FIN, GBR, and TSI, whereas in IBS was not (S1 Fig A); In the second round, the myopia genetic risk increased as latitude decreased in FIN, GBR, IBS, and TSI (Fig 1B, S1B and S1C Fig).

### Detection of selective sweep near *RHO* region

PBS was calculated in three populations to detect selection signals in myopia-associated loci. FIN was found to be the least susceptible to myopia among EUR populations, TSI was found the most susceptible, and YRI was the outgroup. The top 99.9% signals were retained as candidate loci, which were categorized by *P* threshold in the myopia GWAS summary statistics[29], including 159 variants at *P* threshold 5.0 × 10^−3^ and 24 variants at *P* threshold 5.0 × 10^−8^. 21 and 9 nearest genes were obtained from these two sets of variants based on the NCBI database, respectively (S1 and S2 Tables). Of all the top signals in the two different sets, *RHO* was the only gene that was shown to participate in the response to light environments (Fig 2A).

**Fig 2.**
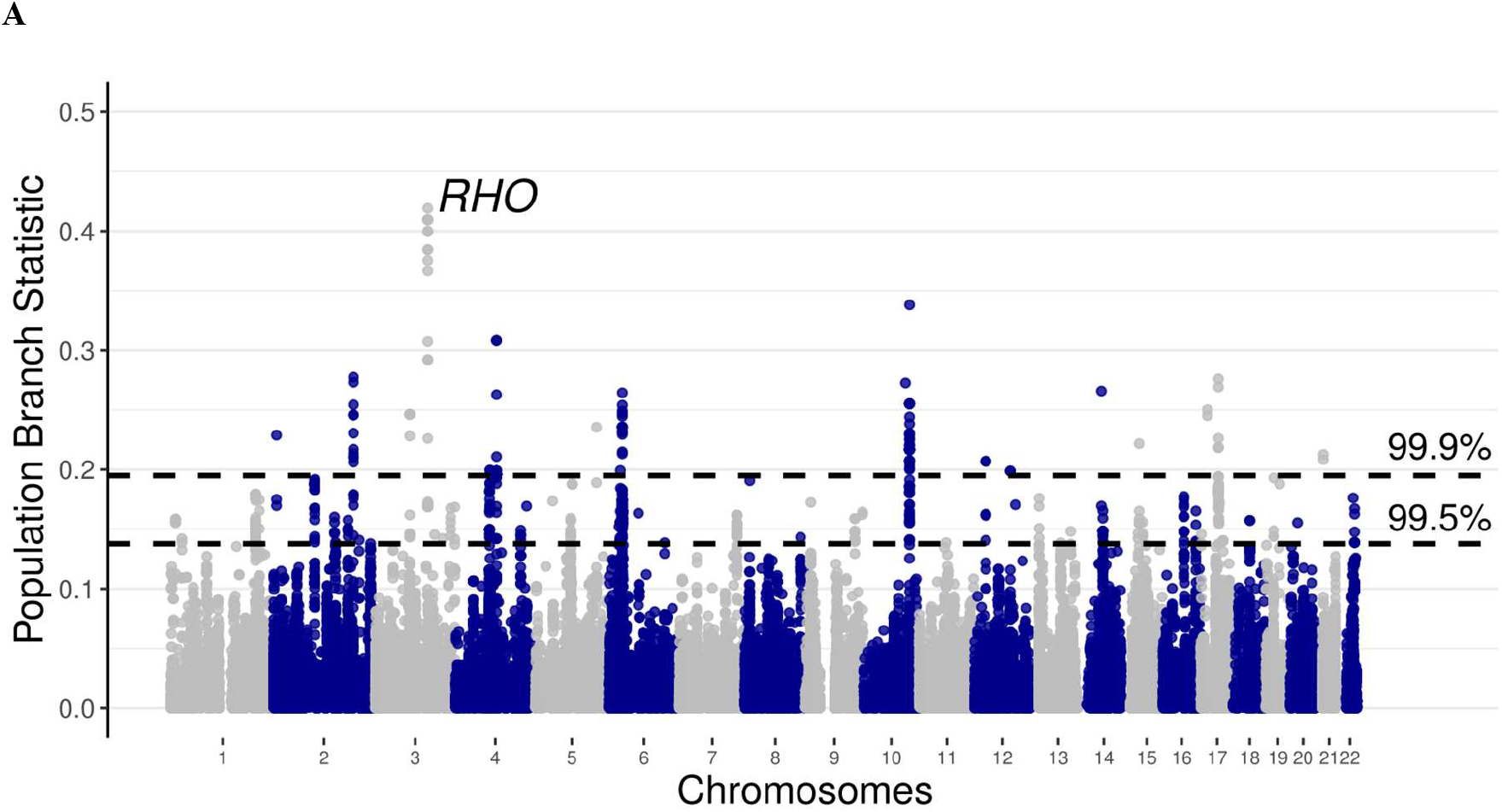
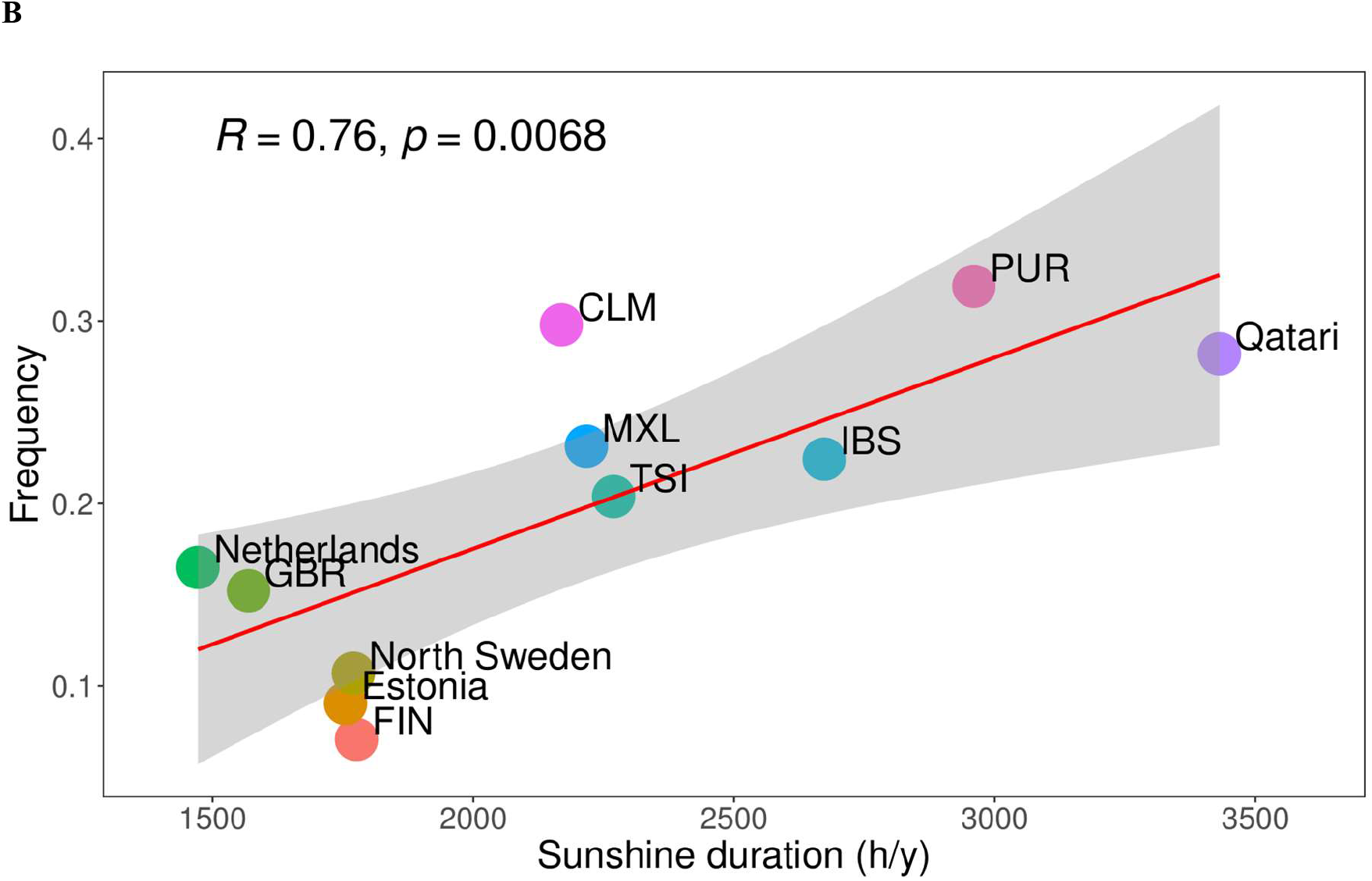
Population Branch Statistics of myopia-associated loci and correlation between identified allele frequencies and sunshine duration. (A) 159,190 SNPs from the PRS-scored variant sets were filtered by *P* < 5.0 × 10^−3^, and the 99.5^th^ and 99.9^th^ percentiles of population branch statistic (PBS) values are shown as black dashed horizontal lines. (B) Correlation between rs2855558 G frequency and sunshine duration of population corresponding city. Sunshine duration data (hours per year) are from the Hong Kong Observatory, allele frequencies are from NCBI, and target data. FIN: Finnish in Finland; GBR: British from England and Scotland; TSI: Toscani in Italia; IBS: Iberian populations in Spain; CLM: Colombian in Medellín, Colombia; MXL: Mexican Ancestry in Los Angeles CA the USA; PUR: Puerto Rican in Puerto Rico.

The five identified variants (rs2625953, rs2625954, rs2625955, rs7984, and rs2855558) near *RHO* were in tight linkage disequilibrium (LD, *r*^*2*^=1) in FIN. To verify whether this LD block was a consequence of selective sweep, we computed *nS*_*L*_[34] and XP*-nS*_*L*_[35] using Selscan 2.0[37] (https://www.github.com/szpiech/selscan). These variants showed a selective sweep signature in FIN, but not in YRI (S2 Fig). Of the five variants, rs2855558 was marked as critical for being top in XP-*nS*_*L*_.

The rs2855558 allele frequency varies worldwide (Table 1). As we found a correlation between myopia genetic risk and latitude, we tested the correlation between rs2855558 allele frequency and the sunshine duration among the worldwide populations. A correlation between the rs2855558 allele frequency and local sunshine duration was observed (*R*=0.19, *P*=0.34, for all 27 populations; *R*=0.61, *P*=0.011, for 16 populations of EUR, AMR, and SAS; Table 1), and was more significant in a subset of European ancestry populations (EUR, AMR, *R*=0.76, *P*=0.0069; Fig 2B, Table 1).

### Diversity of selective pressure of previously identified myopia genes

As this study focused on light adaptation, genes affecting retinal structure and function could have a higher chance to be involved in selection, such as *RHO* which is mainly expressed in rod cells. The *nS*_*L*_ of genome-wide significant myopia-associated loci[29] was calculated in FIN, and genes expressed in retinal layers were categorized based on previous expression analysis[29,30]: 333 genes not directly related to retinal layers, and the 44 genes expressed in retinal cells were further sorted as 35 common refractive error genes and 9 syndromic myopia genes (Fig 3A).

**Fig 3.**
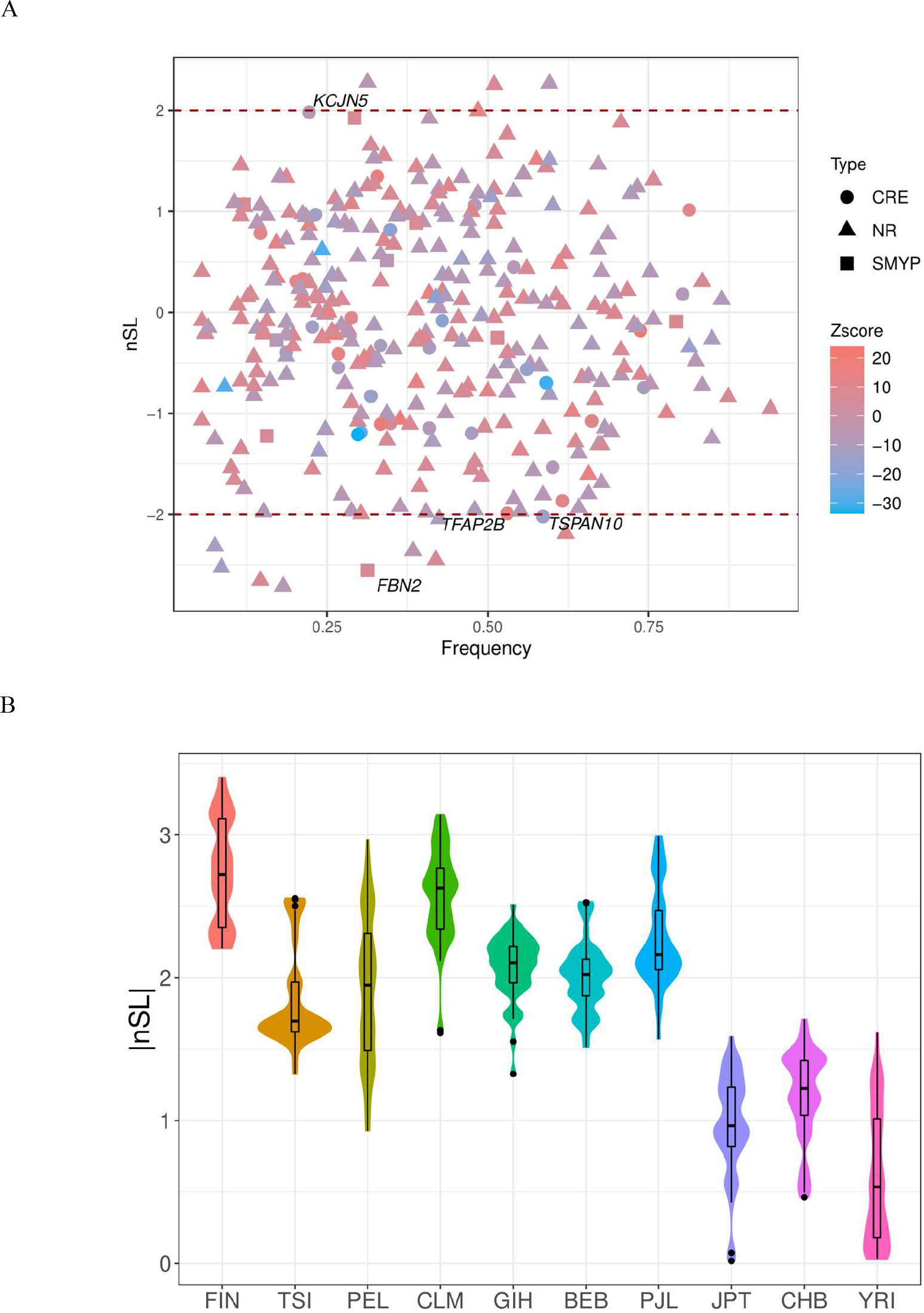
*nS*_*L*_ distribution of myopia-associated loci in FIN and |*nS*_*L*_| distribution of 48 SNPs in LD with rs2855558 in 1KGP populations. (A) *nS*_*L*_ distribution of 333 myopia-associated loci in FIN. *nS*_*L*_ was calculated by Selscan 2.0[37]. Allele frequencies and effect size were from base data[29], loci expressed in retinal layers were categorized based on annotation[29,30], NR: not directly related to retinal layers; CRE: common refractive error genes expressed in retinal layers; SMYP: syndromic myopia genes expressed in retinal layers. (B) |*nS*_*L*_| distribution of 48 SNPs in LD with rs2855558 in 1KGP populations, these SNPs were in LD (*r*^*2*^>0.8) in FIN. FIN: Finnish in Finland; TSI: Toscani in Italia; PEL: Peruvian from Lima, Peru; CLM: Colombian from Medellian, Colombia; GIH: Gujarati Indian from Houston, Texas; BEB: Bengali from Bangladesh; PJL: Punjabi from Lahore, Pakistan; JPT: Japanese in Tokyo, Japan; CHB: Han Chinese in Beijing, China; YRI: Yoruba in Ibadan, Nigeria.

Among genes those were under strong selection (|*nS*_*L*_| ∼ 2 or larger), *FBN2* (*P* = 8.63 × 10^−11^ for rs6860901) was the only syndromic myopia gene; *KCNJ5* (*P* = 1.17 × 10^−17^ for rs10219187), *TAFP2B* (*P* = 4.19 × 10^−37^ for rs2076309), and *TSPAN10* (*P* = 2.22 ×10 ^−50^ for rs9747347) were common refractive error genes[29].

From the combination of effect size and *nS*_*L*_ of the corresponding allele, we could infer the selection indication that if both effect size and *nS*_*L*_ were positive or negative, selection favored hyperopia; if one was negative and the other was positive, selection favored myopia (S3 Table). Since myopia is polygenic, no consistent tendency was observed, while selection favored myopia in FIN for *FBN2* (rs6860901 C) and *TSPAN10* (rs9747347 T).

rs9747347 was also significantly associated with hair color (*P* = 2.94 ×10 ^−21^) in that ancestral allele T refers to red hair and derived allele C to brown and black hair^38^. Considering its significant association with both hair color and myopia, meanwhile selection favored myopia at this locus, we presumed the selective pressure in myopia-associated loci could originate from adaptation events involved in light environment rather than those associated with myopia itself.

### Haplotype pattern and frequency distribution reveals the ethnic specification

The LD block of rs2855558 spanned ∼166.7kb range of chromosome 3 and contained a total of 48 biallelic variants with high LD status (*r*^*2*^>0.8) in FIN. We compared the intensity of selective sweeps across populations in the LD block using |*nSL*|, a clear gap between the AFR-EAS and EUR-AMR-SAS was observed, and the |*nSL*| distribution also supported selective neutrality in the EAS (Fig 3B). The age of the derived rs2855558 A allele was estimated to be no earlier than 20,000 years ago in EUR[39], well after the European and East Asian divergence 22,000–25,000 years ago[40,41,42]; hence, this gap might be due to ancestry divergence.

We also compared the haplotypes of the LD blocks of *TSPAN10* (rs974347) and *SLC24A5* (rs1426654) extracted from a previous study[43] (S4 Fig), which are representative of pigmentation. The *SLC24A5* and *TSPAN10* haplotypes presented similar geographical distribution to *RHO*, indicating that the *RHO* selective pressure was comparable to pigmentation, which was mainly due to adaptation to light environment.

## Discussion

This study aimed to understand the diversity in the genetic background of myopia from an evolutionary perspective, and provide evidence of local adaptation to the sunlight environment in the retina. The protective effect of bright sunlight against myopia progression in humans6-10 and animal models of chickens, guinea pigs, tree shrews, mice, and some nonhuman primates[44,45,46] implies that adaptation to sunlight conditions in the eyes is long known, conserved, and remains active. In this case, the evolutionary history of the genetic background of myopia highly conforms to traits shaped by light-related adaptation[17], such as pigmentation, and eventually played a role in myopia prevalence divergence.

As many genes, such as oculocutaneous albinism *(OCA)* and *TSPAN10*, involved in pigmentation have also been significantly associated with refractive error[29,30], we propose that a proportion of diversity in the genetic background of myopia, especially moderate myopia, as well as iris color and hair color, might be by-products of selection on skin pigmentation[47]. For instance, a less ultraviolet radiation environment at high latitude regions favored skin depigmentation[47], resulting in a light-colored iris, which could affect the genetic background of myopia by allowing more light onto the retina and consequently reflected on myopia progression[48,49,50].

PRS differentiation in EUR revealed an overall tendency of the genetic background of myopia (Fig 1B, S1 Fig). The average PRS of the EUR populations decreased by latitude in both two settings in the second PRS round, one with the 46,423 common variants which only accounted for less than 20% of each EUR population’s scored variants, and the other with the full sets of 569,973 scored variants (Fig 1B). The average PRS of one cohort based on the same set of variants represents the average allele frequencies (n_i_ is the number of effect alleles i; PT is the *P* threshold; β_i_ is the effect size of effect allele i; G_i,j_ is the genotype of effect allele i of individual j, counted by 0, 1, or 2; N is the number of effect alleles).

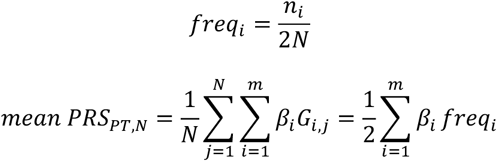

Thus, PRS differentiation indicates frequency spectrum shifts of myopia-associated alleles in EUR, which may correlate with sunshine duration.

Widespread commercial indoor lighting is merely 100 years old. With modern lifestyle becoming indoor-centric, the indoor illuminance (100–500 lx) can hardly match any shaded or cloudy outdoor environments (10,000 to 40,000 lx), whereas clear sunny days are brighter than cloudy days by folds (∼130,000 lx)[51]. Therefore, through adaptation, those consistently living in places of reduced bright sunny hours could have a higher chance of being less susceptible to myopia due to light environment changes, such as Norther Europeans.

Interestingly, the frequencies of derived rs2855558 (*RHO*) A allele in EAS were ∼40%, whereas those of rs1426654 (*SLC24A5*) were 0–3%, and the frequency distributions in AFR, EUR, and SAS were similar between the two alleles (S4 Fig). As we found selective neutrality of rs2855558 in EAS, and the age of its derived alllel was similar to the ages of onset of selective sweeps in *SLC24A5* (rs1426654)[52]. The high proportion of derived rs2855558 A alleles in EAS may be either inherited from ancestors before the divergence of populations from East Asian countries and Europe, or East Asia underwent great genetic drift after the divergence[53], or both. The selective neutrality of rs2855558 in EAS and AFR also explained the robusted correlation between rs2855558 allele frequencies and sunshine duration in a subset of European ancestry populations (Fig 2B).

We also noticed that the frequencies of derived rs1426654 and rs9747347 (*TSPAN10*) alleles in PEL were lower than those in CLM and PUR in AMR populations, while the frequencies were higher in PEL for rs2855558. AMR ancestors migrated from East and North Asia[54], and PEL has been previously reported to share the least alleles with EUR among AMR[36]. The origin of this allele frequency differentiation in AMR was verified by haplotype comparisons (S4 Fig), that haplotypes including derived alleles in EAS were different from PEL in *RHO*; thus, the allele frequency deviation of the three loci in PEL may be due to admixture state variation with EUR instead of from Asian ancestors. Besides, |*nS*_*L*_| of *RHO* variants in PEL were found under strong selection (Fig 3B). Although lacking uniform resources of sunshine duration data, Lima of Peru is well known for its cloudy and foggy weather due to the Humboldt Current. Hence, local adaptation could have also played a role in the elevated rs2855558 allele frequency in PEL.

The validation of the myopia genetic background divergence between EAS and EUR remains difficult because major large-scale myopia GWAS were from European populations[4,29,30], and studies of comparable power in East Asians are absent. Particularly, the base data were more representative of the genetic risk for common myopia than for high myopia in the meta-GWAS[29].

In a Singaporean study[55], the same summary statistics[29] were deduced to predict high myopia (spherical equivalent≤ -5.00D) in East Asians, but the prediction was not performed well for moderate myopia (−5.00 D <spherical equivalent ≤ −3.00 D). The main feature of their approach was filtering PRS scoring SNPs by minor allele frequency (MAF) no less than 0.01 in Chinese children. The authors found that by adding PRS into predictors, the difference in prediction performance between high and moderate myopia was enlarged, and they also observed a small improvement in prediction performance only for moderate myopia when the PRS was generated using 683,970 HapMap3 SNPs than using their approach. As SNPs extracted from this approach were prone to represent the genetic risk for high myopia, combined with our findings, we infer that the genetic background of high myopia is comparable between EAS and EUR, while the divergence of the genetic background of common myopia might be underestimated.

This study applied a two-round PRS strategy to compare the general genetic risk among EUR populations, which has the potential to avoid noise from admixture and boost accuracy for cross-population comparison. In the first-round PRS results, the IBS bias represented more intense LD pattern variations because the default variant sets from clumping in PLINK would drastically shift in populations of different LD patterns, resulting in an unpredictable bias in PRS. Noises will be amplified even within a tolerable genetic distance, that is, between UKBB and IBS.

Although *RHO* was found under strong selection in myopia-associated loci, decisive evidence was missing that selection on myopia or other phenotypes related to light environment was behind the allelic diversity. Depigmentation could also ease the environmental impact; iris of lighter colors can let more light getting into the retina, so the possibility that the high frequencies of derived alleles in *RHO* can increase the expression of *RHO* may not be essential in insufficient sunshine regions in Europe. The correlation between iris color and myopia[37,38] may also be from the divergence of genetic background that pigmentation was the feature instead of causation. But our findings revealed the tip of the iceberg of understanding how adaptation could shape the diversity of the genetic background of myopia.

In summary, myopia susceptibility correlated with sunshine duration in the EUR, and selective signatures were detected in the *RHO*, indicating that local adaptation played a role in the formation of the myopia genetic background. The diversity in the genetic background of myopia may originate from the selection of pigmentation and interaction with depigmented traits, such as light-colored eyes, which could potentially allow more light to enter the retinal layers. EAS and EUR had different adaptation histories of myopia, possibly resulting in underestimated divergence in the genetic background of moderate myopia. Despite these findings, the true driver of diversity of *RHO* remains uncertain, as well as whether and how skin pigmentation played a role in the diversification of the myopia genetic background in EAS and PEL. The next step requires the latest myopia prevalence records and uniform sunshine duration records to estimate the effect size of environmental impact; further validation of the genetic background divergence between EAS and EUR is needed based on East Asian myopia GWAS results of equivalent power to the base data.

## Materials and Methods

### Quality control of the base data and target data

Base data[29] were extracted from previous myopia meta-GWAS of European ancestry (ftp://twinr-ftp.kcl.ac.uk/Refractive_Error_MetaAnalysis_2020), the raw summary statistics contains 7,582,481 variants. The following quality control criteria were used to remove variants: MAF less than 0.01, duplicated variants, ambiguous SNPs, and mismatched SNPs. Ultimately, 6,404,424 variants were identified.

Genotype data of 80,855,802 variants produced by whole-genome resequencing of the 1000 Genomes Project phase 3 populations were used as the target data[36]. VCF files were downloaded from open access of IGSR (ftp://ftp.1000genomes.ebi.ac.uk/vol1/ftp/release/20130502/). Target data were filtered by successful genotyping rate > 0.99, sample missingness < 0.1, Hardy-Weinberg equilibrium *P* >1 × 10^−6^, heterozygosity within three standard deviations of the mean, and MAF > 0.01. After quality control, 9,998,585 variants were retained for FIN (N=96), 9,963,795 for GBR (N=91), 10,476,106 for IBS (N=100), and 9,365,823 for TSI (N=107).

### Polygenic Risk Scores

To compare the genetic risk of myopia among EUR populations, PRS[32] was computed using PLINK v1.9 (https://www.cog-genomics.org/plink/1.9) for 1KGP EUR populations, FIN, GBR, IBS, and TSI.

Since the target data[39] comprise abundant rare variants (0.01<MAF<0.05), most of which were common variants and weakly associated with myopia, more restricted *P* threshold sets were considered for more significant associations, for example, *P* < 5.0 × 10^−5^.

*P*-value-based variant clumping was performed by LD *r*^*2*^ < 0.1 and a physical distance threshold of 250 kb. PRS was calculated per individual at eight *P* thresholds (0.1, 0.05, 0.01, 5 × 10^−3^, 5.0 × 10^−5^, 5.0 × 10^−8^, 5.0 × 10^−12^ and 5.0 × 10^−20^).

Effect sizes were calculated from the Z-score in the base data using the following equation ^56^, where *f* is the effect allele frequency, and *N* is the corresponding weight (effective population size).

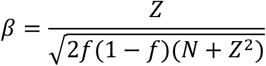

To avoid over-fitting for GBR and random sampling bias during variant clumping due to general genetic distance (e.g., FIN and IBS), PRS was calculated for two rounds: the first round followed the online method[57], scored variant numbers were 216,361 in FIN, 245,340 in GBR, 251,701 in TSI, and 262,336 in IBS; the second round combined 46,423 mutual variants in the first round, and the full set of 569,973 scored variants in the first round was extracted for PRS for validation. The final PRS was uniformed by percentile, as the maximum PRS in each set was equal to 1.

Although phenotypes are lacking to verify the variance proportion explained by each PRS setting, the population risk distribution still reveals the general genetic background differences; nevertheless, the comparison is highly restricted within European ancestries.

### Detection of selective signals and LD haplotype pattern comparison

PBSs^33^ were calculated using myopia-associated variants extracted from the base data and genotypes from the target data. Since FIN and TSI were the most differentiated in PRS results, PBS was calculated between the two populations, and YRI was calculated as the out-group. PBS was calculated at two sets of the PRS-scored variants, *P* threshold at 5.0× 10^−3^ and 5.0 × 10^−8^, which refer to suggestive significant and genome-wide significance, respectively[58].

PBS only revealed the relative selection signal among target populations; to filter the signal under strong selection, *nS*_*L*_[34] and XP*-nS*_*L*_[35] were further computed using Selscan 2.0[37]. Computing variants were limited to phased and biallelic loci, autosome genotypes of FIN, TSI, PEL, CLM, GIH, BEB, PJL, JPT, CHB, and YRI were inputted after QC, and the results were normalized before further analyses. To avoid bias from admixture or genetic drift, at least two populations were included for each superpopulation, except for the AFR.

Variants (S2 Table) in LD with rs2855558 (*r*^*2*^>0.8) were retrieved for haplotype comparisons (S4 Fig). FIN, TSI, PEL, CLM, GIH, BEB, JPT, CHB, and YRI were chosen to represent each superpopulation in 1KGP3, and haplotypes of 2*N*=1778 were compared by population and ancestral-derived alleles. The haplotypes of *TSPAN10* (rs9747347) were constructed using the same approach (S4 Fig). Haplotypes of *SLC24A5* (rs1426654) were built based on common SNPs between 1KGP and a previous study^43^ due to the poor LD estimation performance of rs1426654 in FIN and TSI.

### Correlation between allele frequency and sunshine duration

Sunshine duration data (hours/year) were retrieved from the Hong Kong Observatory (https://www.hko.gov.hk), and a unified data source guaranteed comparability. Ancestral allele frequencies of rs2855558 (G) were obtained from the base data and NCBI database. The correlation between ancestral allele frequency of rs2855558 (G) and sunshine duration was conducted by three groups: all 27 populations had sunshine hours records; 16 populations excluded AFR and East Asian; and 10 populations including northern Europeans, EUR, and AMR (PEL was excluded for no unified sunshine duration record).

## Data, Materials, and Software Availability

Genotype data, GWAS summary statistics, and softwares used in this study are publicly available from the original data providers.

## Acknowledgments

This study was partly supported by JSPS KAKENHI Grant number J21411010, a Grants-in-Aid for Scientific Research program provided by the Japan Society for the Promotion of Science.

## Notes

### Competing Interest Statement

The authors have declared no competing interest.

